# Transient mRNA-LNP Delivery of the Tardigrade Protein Dsup Protects Airway Epithelium from Bleomycin-Induced Injury

**DOI:** 10.64898/2026.11.25.769277

**Authors:** Ian C. Sutton, Pascal Cuhat, Alexander Watson, Christine Wohlford-Lenane, Erin J. Elizalde, Cory Fines, Ameya R. Kirtane, Ashley L. Cooney, Peter J Szachowicz, James D. Byrne

## Abstract

Bleomycin is an essential component of chemotherapy regimens used for Hodgkin lymphoma and germ cell tumors, but its clinical utility is restricted by pulmonary toxicity that forces dose reduction or discontinuation in around 10% of patients. There are currently no approved strategies that prophylactically protect the lungs, representing a significant unmet clinical need. Bleomycin generates reactive oxygen species (ROS) within the alveolar epithelium, causing alveolar cell death and senescence and driving maladaptive crosstalk with stromal cells. Tardigrades are extremophilic organisms that can survive high levels of ROS. This is in part through expression of the tardigrade-specific Damage suppressor (Dsup) protein, which interacts with chromatin and shields the tardigrade genome from ROS. Induction of Dsup expression has been shown to protect human epithelia from radiation-induced injury, and because radiation and bleomycin share ROS generation as their principal cytotoxic mechanism, we hypothesized that this protection may extend to chemoprotection within the alveolar epithelium. We therefore asked whether transient, non-integrating Dsup expression delivered by messenger RNA lipid nanoparticles (mRNA-LNPs) could protect lung epithelium during the window of drug exposure. We employed an LNP formulation that retained its physicochemical properties after nebulization and produced lung-restricted reporter expression after intratracheal administration in mice. Additionally, the same formulation successfully transfected human and porcine air-liquid-interface cultures. Dsup-GFP mRNA transfection produced nuclear-restricted expression in A549 and primary human small airway epithelial cells (HSAECs) that peaked at 24 hours and returned to background by day 5, with no effect on proliferation before or after clearance. Central to its potential chemoprotective role, Dsup mRNA LNP pretreatment significantly reduced bleomycin-induced DNA damage and senescence in both cell types. These results establish transient mRNA-delivered Dsup as a feasible epithelial chemoprotective strategy and define a delivery platform compatible with inhaled administration.

Figure 1.
Graphical abstract
The tardigrade Damage Suppressor protein (Dsup) shields tardigrade chromatin from reactive oxygen species-mediated DNA damage. Here, Dsup is encoded as synthetic mRNA and encapsulated in lipid nanoparticles formulated for pulmonary delivery, enabling transient, non-integrating Dsup expression in the airway epithelium. Inhaled Dsup mRNA nanoparticle pretreatment is proposed as a chemoprotective strategy to prevent bleomycin-induced lung toxicity.

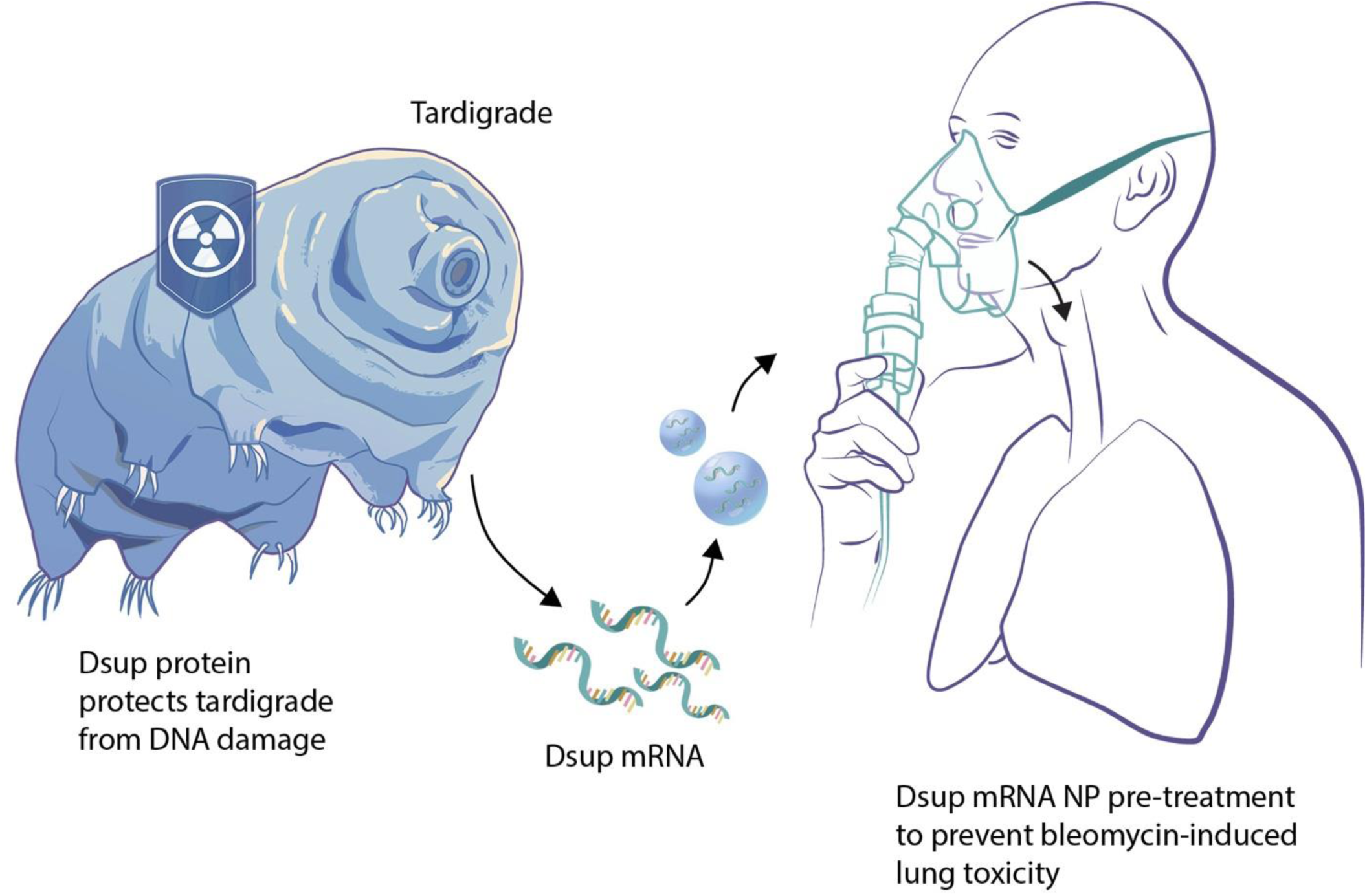

## INTRODUCTION

Bleomycin remains a component of curative-intent chemotherapy for Hodgkin lymphoma and germ cell tumors, where it contributes meaningfully to regimens used in young patients with excellent long-term survival (*1–3*). Its use is constrained by pulmonary toxicity, which occurs in approximately 10% of treated patients, with reported incidence ranging from 7% to 36% depending on cumulative dose, age, renal function, and concurrent oxygen exposure (*3, 4*). Bleomycin-induced pneumonitis progresses to fibrosis in a subset of patients which is irreversible and fatal (*3, 5*). Because toxicity is cumulative and unpredictable at the individual level, clinical management is limited to dose capping, surveillance, and discontinuation (*6*). These strategies trade oncologic efficacy for pulmonary safety. Therefore, there is an urgent need to develop a novel approach to prophylactically protect the lung and ensure ability of clinicians to optimally dose bleomycin.

The cellular mechanism of this toxicity is well characterized. Bleomycin associates with divalent cations such as ferrous ions (Fe^2+^) and interacts with molecular oxygen and water to generate reactive oxygen species that produce single- and double-strand DNA breaks (*7, 8*). This places the alveolus, the body’s highest-oxygen compartment, at greatest risk. Additionally, bleomycin hydrolase, the enzyme that inactivates the drug, is expressed at low levels in lung (*9, 10*). Critically, the injured cells include type II alveolar cells, which are the progenitor pool of the alveolar epithelium. Injury to these cells reduces the capacity for repair and is alone sufficient to drive fibrosis in mice (*11, 12*). Epithelial DNA damage drives apoptosis and senescence, and senescent epithelium activates adjacent fibroblasts through the senescence-associated secretory phenotype (*13–17*). This initiates the pro-fibrotic cascade that culminates in matrix deposition and remodeling, which are the hallmarks of pulmonary fibrosis (*14, 18*). Protection of the epithelium at the level of DNA integrity is mechanistically upstream of fibrosis, yet no approved agent acts there. One of the only approved free radical-scavenging cytoprotectants, amifostine, is indicated for cisplatin-induced nephrotoxicity and radiation xerostomia rather than pulmonary injury (*19*). Its label explicitly advises against use where chemotherapy is curative, naming germ cell malignancies, because systemic cytoprotection may interfere with antitumor activity (*20*). Systemic administration and strong side effect profile (dose-limiting hypotension and emesis) have further restricted its adoption (*19, 21*).

Tardigrades are extremophilic organisms that can survive ionizing radiation doses lethal to most other organisms, in part due to expression of the damage-suppressor protein (Dsup) (*22, 23*). Dsup is a nucleosome-binding protein that physically shields chromatin from hydroxyl radical-mediated DNA damage (*24, 25*). Our lab has recently shown that induction of Dsup expression in oral and intestinal epithelia can reduce radiation-induced DNA damage, establishing that the protective mechanism transfers across species and does not require tardigrade-specific cofactors (*26*). Since bleomycin and ionizing radiation converge on ROS-mediated DNA damage as their principal cytotoxic mechanism, this protection should in principle extend to bleomycin-induced injury of the alveolar epithelium. Dsup is therefore an appealing candidate for protecting normal tissue against bleomycin-induced pulmonary toxicity, when timing is known in advance, as it is for almost all scheduled chemotherapy infusions.

Lipid nanoparticle (LNP)-mediated mRNA delivery is well suited to this requirement. mRNA-LNPs produce protein without genomic integration, and their lipid composition can be tuned to favor pulmonary deposition and epithelial uptake (*27–29*). Chemotherapy is administered on a known schedule, so protection is needed only during the window of drug exposure. An mRNA pulse matches that requirement directly, as Dsup is present while the epithelium is at risk and cleared afterward, leaving normal tissue unmodified.

Here we report a nebulization-stable LNP formulation that achieves lung-restricted mRNA expression in mice and transfects human and porcine airway epithelial cultures at the air-liquid interface. We show that Dsup-GFP mRNA delivered by this platform localizes to the nucleus, is cleared within five days, and does not negatively alter cell proliferation. We then show that Dsup pretreatment reduces bleomycin-induced DNA double-strand breaks, total DNA damage, and cellular senescence in both an immortalized alveolar epithelial cell line and primary human small airway epithelial cells. Together these data establish transient mRNA-delivered Dsup as a novel strategy for alveolar epithelial chemoprotection.

## RESULTS

### A nebulization-stable LNP formulation achieves lung-restricted mRNA expression

A chemoprotective agent for bleomycin-induced lung injury must reach the alveolar epithelium and should ideally be deliverable by inhalation. We formulated LNPs containing the ionizable lipid KC2 (35 mol%), DOPE (46.5 mol%), cholesterol (16 mol%), and PEG-lipid (2.5 mol%) and first asked whether they survive aerosolization (Figure 2A). LNPs were constructed with firefly luciferase (FLuc) mRNA, aerosolized using a vibrating mesh nebulizer (volume median diameter 2.5-4.0 µm), and collected directly into a conical tube for analysis. Physicochemical characteristics and encapsulation efficiency were measured pre- and post-nebulization. Nebulization produced no significant change in average particle size (z-average), polydispersity index, zeta potential, or encapsulation efficiency (Figure 2B), indicating that the formulation does not aggregate or destabilize under the shear stress of nebulization.

**Figure 2.**
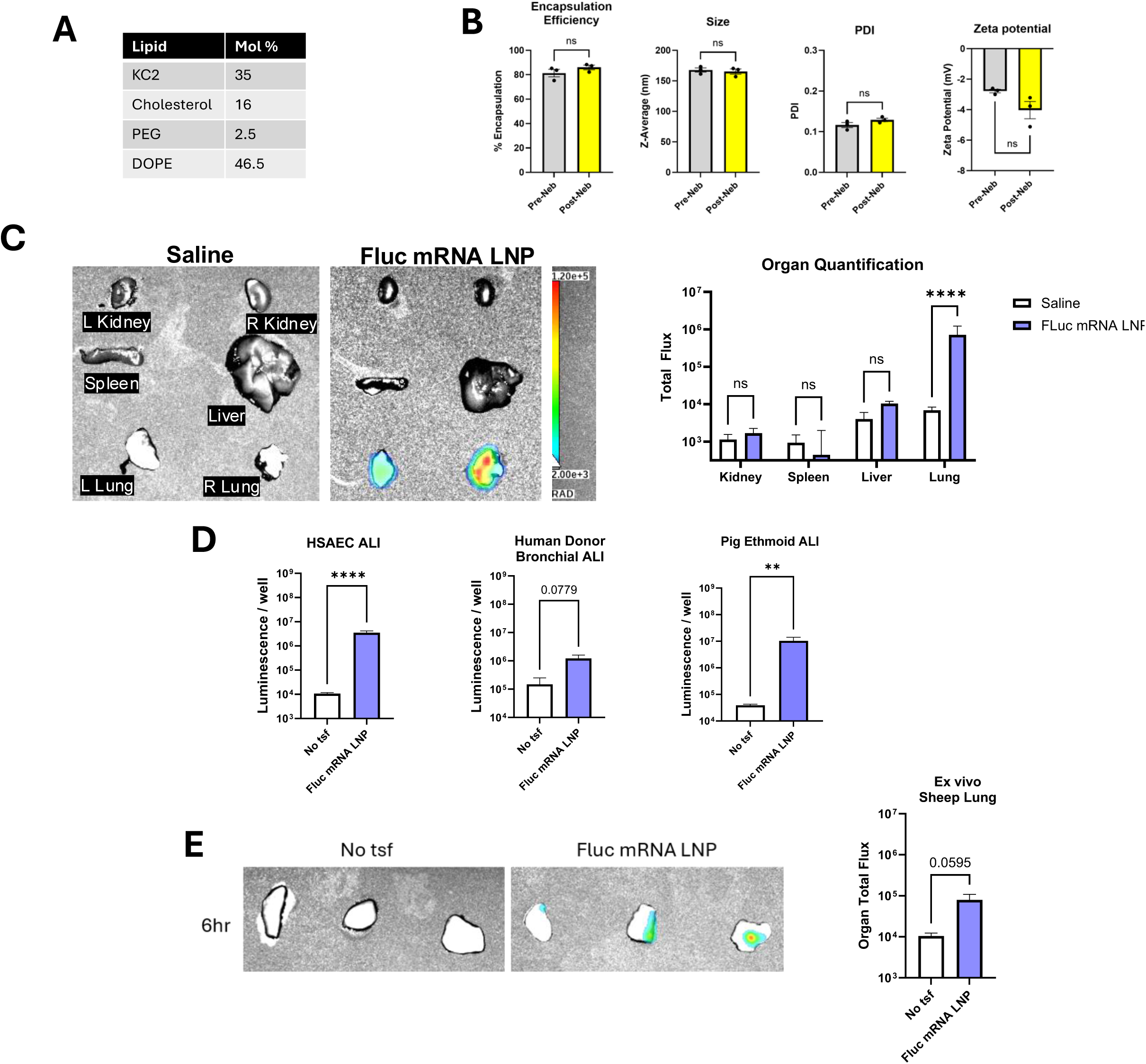
Nebulization-stable LNPs achieve lung-restricted mRNA expression across multiple in vivo, ex vivo, and air-liquid interface models. **(A)** Lipid composition of the LNP formulation (mol %). **(B)** Encapsulation efficiency and physicochemical characterization before and after nebulization. Z-average hydrodynamic diameter, polydispersity index (PDI), and zeta potential were measured by dynamic light scattering. Properties were compared by Welch’s unpaired t-test. **(C)** Representative IVIS images and quantification of organ total flux in mice (n = 5 per group) after intratracheal administration of saline or FLuc mRNA LNP (12.5 µg mRNA), harvested 6 hours post-delivery. Log₁₀-transformed luminescence was compared by two-way ANOVA (treatment × organ) with Šídák’s multiple comparisons test comparing saline versus FLuc mRNA LNP within each organ. **(D)** FLuc mRNA LNP transfection of human small airway epithelial cell (HSAEC), primary human donor bronchial, and pig ethmoid air-liquid interface (ALI) cultures, with luminescence measured 24 hours after apical dosing; No tsf, untransfected. Log₁₀-transformed luminescence was compared by Welch’s unpaired t-test. **(E)** Representative IVIS images and quantification of ex vivo sheep lung slices 6 hours after FLuc mRNA LNP treatment. Log₁₀-transformed luminescence was compared by Welch’s unpaired t-test. Data are mean ± SEM and n = 3 biological replicates except (C), n = 5 mice per group. Exact p values are shown for selected non-significant comparisons. [ns, not significant; **p < 0.01, ****p < 0.0001]

To determine how effectively these particles transfect the lung, nebulized FLuc mRNA LNPs were collected and administered intratracheally to 10-week-old female CD-1 mice. A dose of 12.5 µg FLuc mRNA generated significant reporter expression, approximately a 2-log increase above background, by *ex vivo* bioluminescence imaging at 6 h post-delivery. Because lung delivery does not always equate to lung containment, we also assessed luminescent signal in the liver, spleen, and kidneys of each mouse. Signal was confined to the lungs, with no significant increase in reporter expression in any other organ assessed (Figure 2C). Expression was therefore lung-restricted rather than merely lung-enriched, which is critical for a chemoprotectant designed to protect the lung without interfering in other organ systems where cancer cells may be present.

We next asked whether the platform transfects differentiated airway epithelium from other species and from other levels of the airway. FLuc mRNA LNPs increased bioluminescence signal in HSAEC and pig ethmoid air–liquid interface (ALI) cultures (Figure 2D). In primary human donor bronchial ALI cultures the increase was not statistically significant (p = 0.078), though signal rose by approximately one log, likely reflecting high donor-to-donor variability in transfection efficiency. In *ex vivo* sheep lung, FLuc mRNA LNP treatment produced a signal increase that approached but did not reach significance (p = 0.0595, Figure 2E). Taken together, the formulation transfects differentiated airway epithelium across multiple species and at different points along the airway, with the strongest and most reproducible effects in human small airway and pig ethmoid cultures.

### Dsup-GFP is expressed in the nucleus, cleared within five days, and does not alter proliferation

The therapeutic rationale for Dsup mRNA delivery rests on expression being both correctly localized and genuinely transient. The first model used in this study was A549, a human lung adenocarcinoma-derived line widely used as an accessible *in vitro* model of alveolar type II epithelial cells. Given its malignant origin, all findings required verification in primary cells, for which we used primary human small airway epithelial cells (HSAECs). Both were readily transfected by our LNP formulation carrying mRNA encoding a Dsup-GFP fusion protein or GFP alone as a control. mRNA concentrations used throughout the study were 0.5 µg/mL for A549 and 0.25 µg/mL for HSAECs. To assess localization of the protein encoded by the delivered mRNA, cells were imaged at 24 hours post-transfection by confocal microscopy (Figure 3A). GFP mRNA transfection produced uniform GFP signal throughout the cytoplasm, whereas Dsup-GFP was confined to the nucleus in both cell types, consistent with the nucleosome-binding behavior reported for Dsup in other systems.

**Figure 3.**
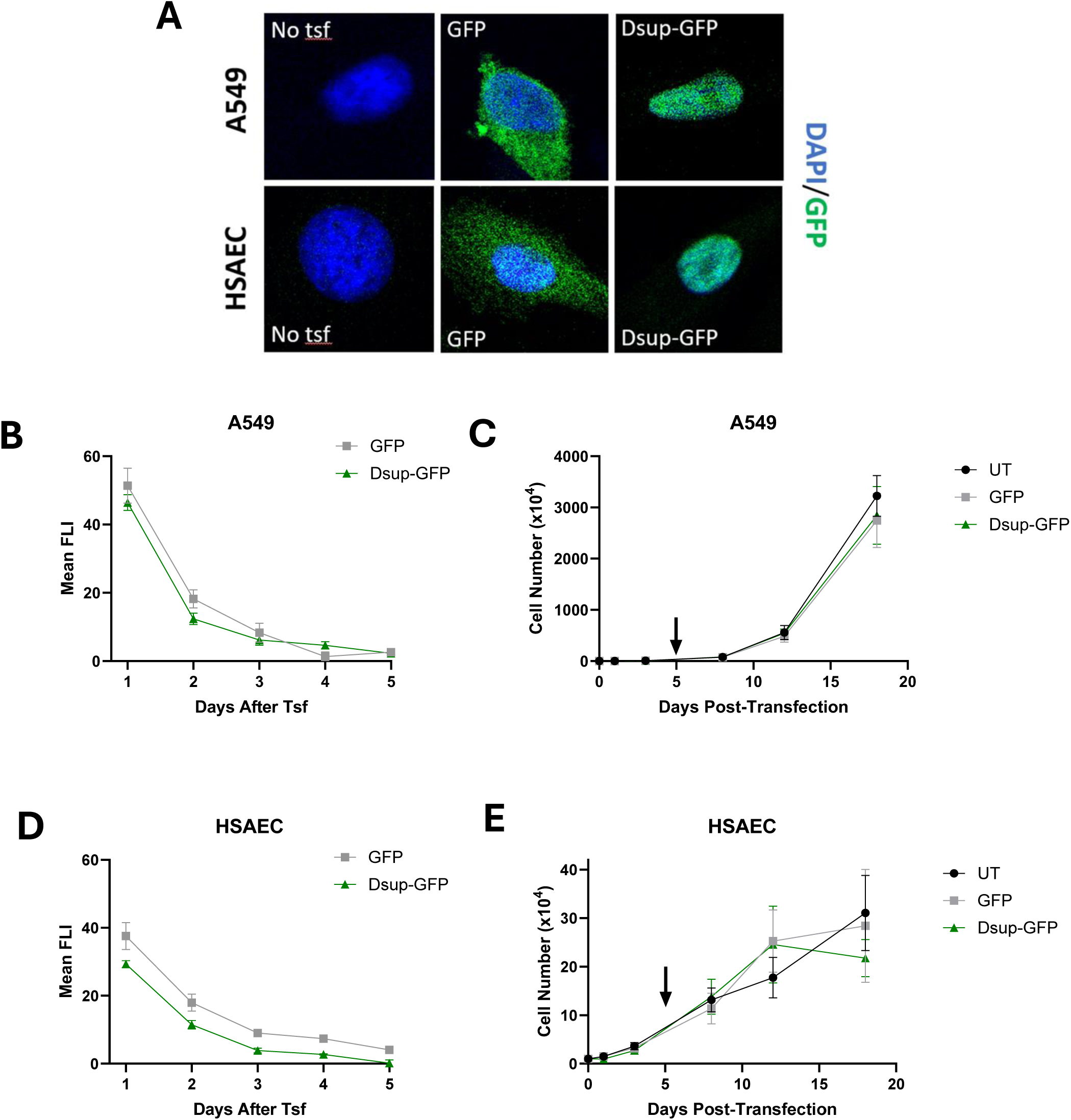
Dsup-GFP expression kinetics parallel those of GFP, and neither construct affects proliferation, in HSAEC and A549 cells following mRNA transfection. **(A)** Representative immunofluorescence images of single A549 and HSAEC cells 24 hours after transfection with GFP or Dsup-GFP mRNA LNP, with untransfected controls (No tsf). Green, GFP; blue, DAPI counterstain. GFP alone distributed throughout the cytoplasm, whereas Dsup-GFP was confined to the nucleus in both cell types. **(B, D)** Mean cell fluorescence intensity (FLI) of GFP and Dsup-GFP in A549 (B) and HSAEC (D) cells over 5 days following transfection (Tsf), measured by fluorescence microscopy. **(C, E)** Cell growth over 18 days post-transfection in A549 (C) and HSAEC (E) cells for untransfected (UT), GFP-, and Dsup-GFP-transfected conditions. Arrow indicates the timepoint at which expression of both constructs had returned to background. Data are mean ± SEM of 3 biological replicates.

Expression kinetics followed the profile expected of an mRNA therapeutic. Mean fluorescence intensity peaked one day after transfection and declined to background by day 5 in both A549 and HSAEC cells (Figure 3B, 3D). Dsup-GFP signal was consistently somewhat lower than GFP at matched timepoints, which is expected for a larger fusion construct and does not affect the kinetics of clearance. Transient expression of a chromatin-binding xenoprotein must also be shown not to perturb basic cell behavior. Neither GFP nor Dsup-GFP altered proliferation relative to untransfected controls over 18 days in A549 cells (Figure 3C). HSAEC growth curves were similarly unaffected, though primary cells showed greater variability at later timepoints (Figure 3E). Growth was tracked well beyond the point of Dsup clearance, so these curves capture delayed consequences of the expression pulse as well as its immediate effects, both relevant to safe clinical translation.

### Dsup pretreatment reduces bleomycin-induced DNA damage

Dsup-mediated protection of epithelial DNA is central to our hypothesized mechanism, so we next tested whether Dsup mRNA LNP pretreatment protects these cells from bleomycin-induced DNA damage. Cells were pretreated with GFP or Dsup mRNA LNPs and exposed to bleomycin at 2 µg/mL for 1 hour. Bleomycin markedly increased DNA double-strand breaks, as measured by γH2AX immunofluorescence staining, and Dsup pretreatment significantly reduced foci burden in A549 cells relative to GFP-pretreated controls (Figure 4A). Dsup expression alone did not alter baseline foci, indicating that the protein does not itself induce a DNA damage response detectable by this assay. HSAECs showed the same pattern (Figure 4B).

**Figure 4.**
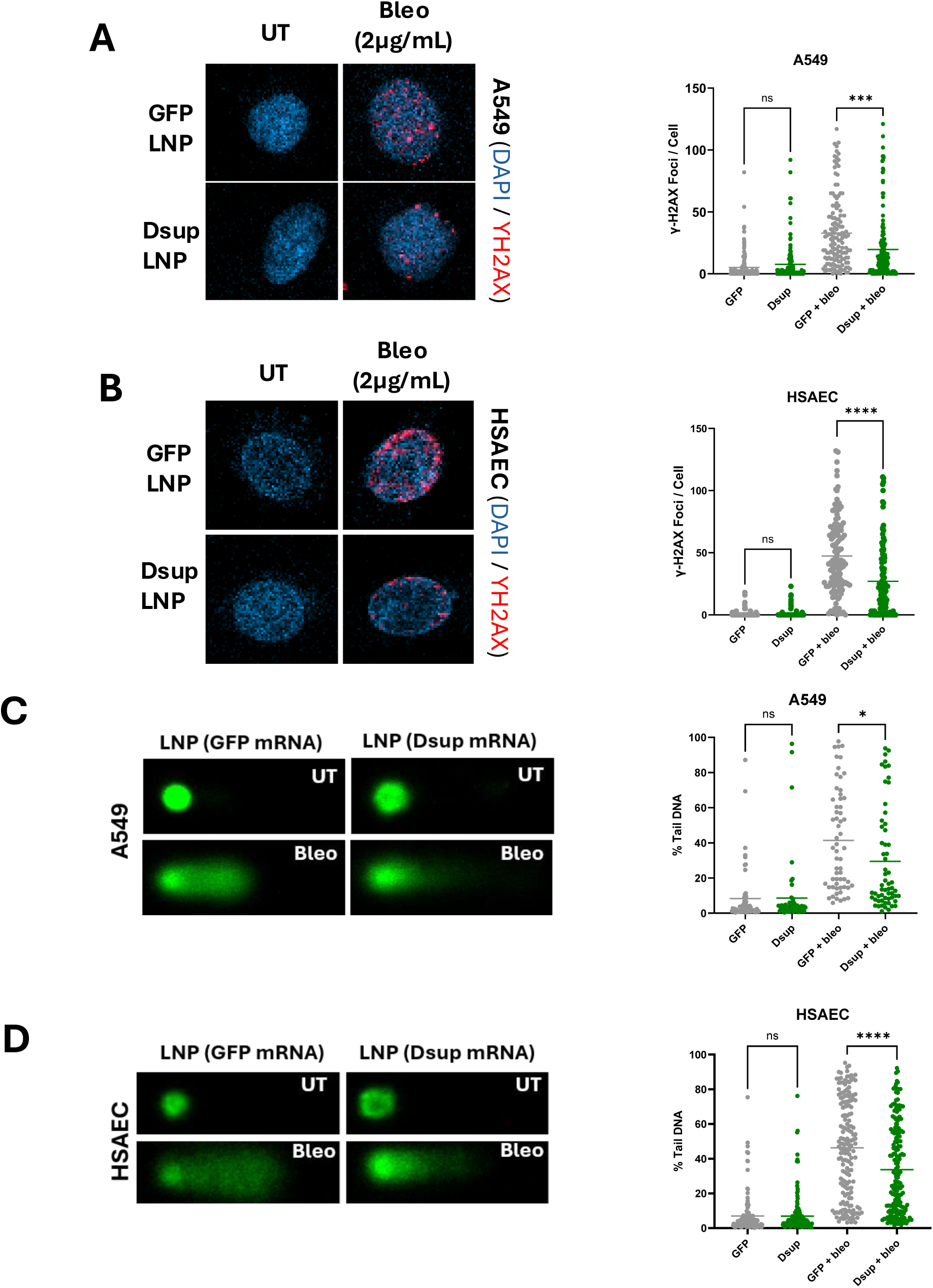
Dsup mRNA-LNP pretreatment reduces bleomycin-induced DNA damage in A549 and primary HSAEC cells. (A, **B)** Representative confocal images of γH2AX immunofluorescence (red) with DAPI counterstain (blue) in A549 (A) and HSAEC (B) cells pretreated with GFP or Dsup mRNA LNP and subsequently left untreated or exposed to bleomycin (2 µg/mL) for 1 hour, fixed immediately after exposure. Quantification shows γH2AX foci per cell. Bleomycin markedly increased foci in GFP-pretreated cells, and Dsup pretreatment significantly reduced foci burden, while Dsup alone did not alter baseline foci. **(C, D)** Representative alkaline comet assay images and quantification of % tail DNA in A549 (C) and HSAEC (D) cells pretreated with GFP or Dsup mRNA LNP and exposed to bleomycin as in (A). Dsup pretreatment significantly reduced % tail DNA in both cell types without affecting baseline damage. Each point represents a single cell, horizontal lines indicate the mean. Data were pooled from 3 independent experiments and compared by one-way ANOVA with Tukey’s post hoc test for multiple comparisons. [ns, not significant; *p < 0.05, ***p < 0.001, ****p < 0.0001]

Because γH2AX reports double-strand breaks specifically, we used alkaline comet assays to capture total DNA damage including single-strand breaks and alkali-labile sites. Dsup mRNA LNP pretreatment significantly reduced percent tail DNA after bleomycin in both A549 cells (Figure 4C) and HSAECs (Figure 4D), with no effect on baseline damage in either cell type. Protection was therefore evident across two mechanistically distinct damage readouts and in both an adenocarcinoma-derived alveolar epithelial line and primary human small airway cells.

### Dsup pretreatment attenuates bleomycin-induced senescence

While DNA damage is the initiating insult to the epithelium, senescence is the state through which damaged epithelium drives fibrosis, via secretion of factors comprising the senescence - associated secretory phenotype. We therefore measured senescence-associated β-galactosidase positivity over 16 days following a 48-hour bleomycin exposure at 0.1 and 1 µg/mL (Figure 5A).

**Figure 5.**
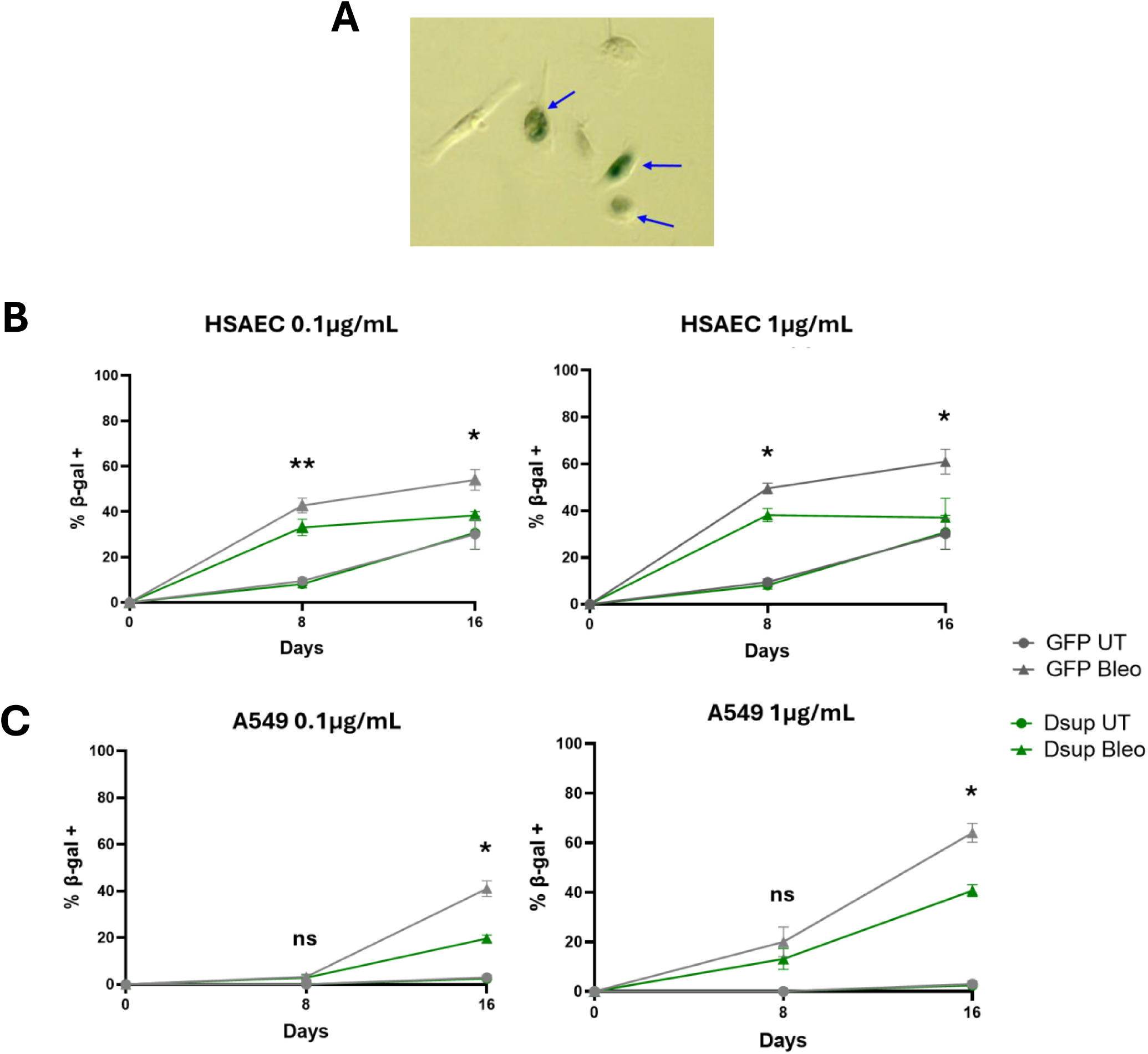
Dsup mRNA LNP pretreatment attenuates bleomycin-induced cellular senescence. **(A)** Representative senescence-associated β-galactosidase (SA-β-gal) staining in HSAECs. Blue arrows indicate SA-β-gal-positive cells. **(B, C)** Percentage of SA-β-gal-positive cells on days 8, and 16 after bleomycin exposure (0.1 or 1 µg/mL for 48 hours) in HSAEC (B) and A549 (C) cells pretreated with GFP or Dsup mRNA LNP. Bleomycin induced progressive senescence in both cell types at both doses, and Dsup pretreatment significantly reduced the senescent fraction. GFP- and Dsup-pretreated bleomycin-exposed groups were compared at days 8 and 16 by two-tailed paired t-tests, matched by independent experiment. Asterisks denote GFP Bleo versus Dsup Bleo comparisons. Data are mean ± SEM, n = 3 biological replicates. [ns, not significant; *p < 0.05; **p < 0.01]

In HSAECs, bleomycin produced a progressive rise in SA-β-gal–positive cells at both doses, reaching approximately 55% and 60% by day 16 at 0.1 and 1 µg/mL respectively. Dsup pretreatment significantly reduced the senescent fraction at both doses and at both day 8 and day 16 (Figure 5B). A549 cells showed lower baseline senescence and a later onset, with significant Dsup-mediated reduction evident at day 16 at both doses (Figure 5C). Dsup expression alone did not increase senescence above GFP controls in either cell type, a necessary safety observation for a protein that binds chromatin globally.

## DISCUSSION

In this study we show that a nebulization-stable lipid nanoparticle formulation delivers Dsup mRNA to lung epithelial cells in a transient, nucleus-restricted fashion. This brief pulse of expression is sufficient to reduce bleomycin-induced DNA damage and senescence in both an alveolar epithelial line and primary human small airway cells. Together these findings establish the proof of concept that the chromatin-protective xenoprotein Dsup can be used therapeutically in the setting of bleomycin-induced pulmonary toxicity.

The transience of expression is a central design feature rather than an incidental property of this strategy. Chemotherapy exposure occurs at a scheduled time, and protection is required only during that window. An mRNA-LNP pulse that peaks within 24 hours and resolves by day 5 matches the duration of the intervention to the duration of the need and leaves no permanent modification of the epithelium behind. This distinguishes the approach from the transgenic and stably integrating systems used in prior demonstrations of Dsup-mediated protection, which commit the cell to indefinite expression of a xenoprotein (*22*).

That distinction raises an important question this study does not address. Dsup binds chromatin nonspecifically across the genome and sustained expression has consequences that differ by system. For example, constitutive expression altered chromatin accessibility in rat cortical neurons and promoted DNA damage (*30*). Additionally, when constitutively expressed in a Drosophila model, Dsup acted as a non-specific transcriptional repressor (*31*). And, most relevant to this study, constitutive Dsup expression in human embryonic kidney cells (HEK293) modulated transcription factor activity and altered expression of genes involved in cellular damage response and repair (*32*). Whether a time-limited mRNA pulse produces comparable perturbation during the period of nuclear occupancy is unknown, as is whether any such change persists once Dsup has cleared. Our proliferation data over 18 days, together with the absence of Dsup-induced senescence or baseline DNA damage, offer functional reassurance that the epithelium is not durably altered. However, these are coarse readouts and direct measurement of chromatin accessibility and transcription across the expression window and after clearance would establish whether duration governs the consequences of Dsup expression. Addressing this concern is fundamental step before Dsup-based therapies are advanced clinically.

The delivery results have implications beyond this specific cargo as well. Lung-restricted expression after airway administration, stability through nebulization, and transfection across human and porcine ALI cultures together indicate a platform that could carry other protective or corrective cargoes to the airway epithelium. The lack of significant expression in liver, spleen, and kidney is particularly relevant for a chemoprotective application, where the entire therapeutic premise depends on protecting normal tissue in one anatomic compartment without conferring protection systemically. With continued advances in the field of LNP-mediated mRNA delivery to the alveolar epithelium, we anticipate that mRNA strategies (including Dsup) will confer even greater chemoprotection with decreased toxicities.

The necessary next step is to test whether the established *in vitro* epithelial protection translates into preserved lung function *in vivo*. Our biodistribution data establish that the platform expresses where it needs to but were generated with a luciferase reporter. Confirming equivalent expression and clearance kinetics for the Dsup construct in the murine lung is a prerequisite for efficacy studies. Those studies follow directly from the readouts used here in our proof-of-concept study. Dsup mRNA LNP pretreatment can be given prior to intratracheal bleomycin, with early assessment of alveolar epithelial DNA damage burden to establish that the protection we observe in culture occurs in the intact lung. This can be followed by assessment at peak fibrosis using histological scoring and pulmonary function testing. Such a design would also address the question our *in vitro* data cannot, namely whether protecting the epithelium is sufficient to prevent the downstream stromal response, since fibrotic burden in an intact lung integrates both. Dose scheduling will require optimization, as the interval between pretreatment and chemotherapy must align the expression peak with the window of drug exposure, and repeated bleomycin cycles may require repeated dosing. Finally, demonstrating that protection is confined to normal lung will require tumor-bearing models in which antitumor efficacy and pulmonary protection are measured in the same animal.

## METHODS

### Cell culture

Human small airway epithelial cells (HSAEC) were obtained from ATCC (PCS-301-010) and maintained in Airway Epithelial Cell Basal Medium (ATCC, PCS-300-030) supplemented with the Bronchial Epithelial Growth Kit (ATCC, PCS-300-040). A549 cells were obtained from ATCC (CCL-185) and cultured in Dulbecco’s Modified Eagle Medium/Nutrient Mixture F-12 with 10% FBS. Primary HSAEC were used at passage 2-5 and A549 were used at passage 10-25. All cell lines were routinely tested for mycoplasma and were maintained at 37°C, 5% CO2.

### LNP formulation, mRNA-LNP transfection, and nebulization of LNPs

GFP, Dsup-GFP fusion, and firefly luciferase (FLuc) mRNAs were synthesized by TriLink. All LNPs were prepared in an identical manner as follows. The individual lipids were dissolved at 10 mg/mL concentration in ethanol and stored at -20° C until used. A lipid working solution was created using the following molar ratio: DLin-KC2-DMA (35%), Cholesterol (16%), PEG-PE (2.5%), and DOPE (46.5%). This lipid mix was added to an equal volume of an aqueous solution of mRNA (1 mg/mL), mixed by pipetting 20 times, and vortexed for 30s. The ionizable lipid (KC2):mRNA weight ratio was maintained at 3.2:1. Following vortex mixing, the LNP solution was placed on ice for 10 minutes to allow for complexation. LNPs were subsequently diluted in either sterile saline for *in vivo* use or cell culture media for *in vitro* transfections. Primary HSAECs and A549 cells were transfected at 0.25 µg/mL and 0.5 µg/mL respectively.

Particle size, zeta potential, and polydispersity index (PDI) were assessed with a Zeta NanoZS device (Malvern). Encapsulation efficiency (EE) was defined using Quant-it RiboGreen Reagent and RNA Assay Kit (Invitrogen, R11490). EE% was calculated using the following equation:

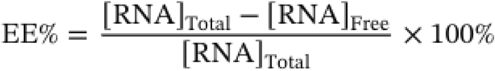

For nebulization stability studies, LNPs were aerosolized using Aeroneb Lab Nebulizer (small volume median diameter, particle size between 2.5µm-4.0µm) into a 15mL conical tube. A seal was created with paraffin between the nebulizer outlet and the top of the 15mL conical tube. The collection tube was placed on ice to allow for condensation of the nebulized LNPs. The nebulizer was allowed to run until LNP solution was fully nebulized and the collection tube was centrifuged at 1,000 rpm for 1 min to bring condensed LNPs on the side of the tube to the bottom. Re-characterization of the LNPs was done immediately post-nebulization for nebulization stability studies.

### Post-transfection cell growth and clearance kinetics

For cell growth assays, A549 and HSAEC were plated in 6 well-plates at a density of 250,000 cells per well. They were transfected at mRNA doses previously specified and were collected 24hr post-transfection. After collection, transfected cells were plated at the following densities for specific collection times:

| HSAEC |  | A549 |  |
|---|---|---|---|
| Growth time | Cell density plated (cells/well) | Growth time | Cell density plated (cells/well) |
| 1 day | 30,000 | 1 day | 10,000 |
| 3 days | 15,000 | 3 days | 10,000 |
| 8 days | 7,500 | 8 days | 1,000 |
| 12 days | 7,500 | 12 days | 500 |
| 18 days | 3,750 | 18 days | 200 |

Cells were counted using the Countess 3 Automated Cell Counter (Invitrogen, AMQAX2000) with trypan blue exclusion. Final cell counts were normalized to reflect initial plating density to construct final growth curves.

To determine the clearance of GFP and Dsup-GFP protein following transfection with their respective mRNAs, cells were plated into 6-well plates and transfected as described above. At 24-hour intervals from 1d-5d post-transfection, live cells were imaged using Evos Fluorescent Microscope and fluorescence intensity (FLI) was measured per cell at each time point. FLI was quantified using Image J. Briefly, mean gray value (average intensity) was taken by outlining cell boundary (GFP) or nucleus (Dsup-GFP). Average background mean gray value was determined by measuring mean gray value in untransfected cells and was subtracted from the post-transfection values.

### Bleomycin treatment

Cells pretreated with GFP or Dsup mRNA-LNP were exposed to bleomycin sulfate (Tocris Bioscience, #7147) at the indicated concentrations prior to downstream analysis. Duration of bleomycin treatment is specified in the individual experiment methods description.

### γ-H2AX immunofluorescence

A549 and HSAEC were plated on sterile 8-well chamber slides and were transfected 24 hours prior to bleomycin challenge at the mRNA doses previously specified. Cells were challenged with bleomycin (2 µg/mL) for 1 hour prior to fixation with 4% paraformaldehyde for 15 minutes. For immunofluorescence staining, cells were permeabilized in a 0.4% Triton-X solution, blocked with 10% FBS solution, and stained with anti-γ-H2AX (phospho S139) antibody (ab22551, Abcam) and DAPI. Corresponding goat anti-mouse 647 secondary antibody (ab150119) was added to conjugate with the anti-γ-H2AX primary antibody. Slides were mounted using Prolong Diamond Antifade with DAPI (Life Technologies, P36962). Images were acquired on Zeiss 990 confocal microscope. Images were taken at the focal plane where maximal discrete foci were observed. Images were uniformly analyzed across all experimental conditions by counting number of discrete foci per cell. 40 cells were analyzed per condition, across 3 biologically independent replicates.

### Alkaline comet assay

A549 and HSAEC were plated at a density of 250,000 cells per well in a 6 -well plate. 24 hours prior to bleomycin challenge, cells were transfected with GFP or Dsup-GFP mRNA at previously specified mRNA doses. Cells were treated with bleomycin (2 µg/mL) for 1 hour, collected from the 6 well-plate, placed into an agarose solution at a density of 1x10^5^ cells/mL, and allowed to solidify on comet slides. Cell lysis and single-cell gel electrophoresis were conducted under alkaline conditions per manufacturer’s instructions (R&D Systems, 4250-050-K). DNA was stained with 1x SYBR gold solution for 30 minutes prior to imaging with Zeiss 990 confocal microscope. Percent tail DNA was quantified using Image J software using the following formula:

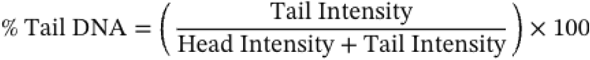

40 cells were analyzed per condition, across 3 biologically independent replicates.

### Senescence-associated β-galactosidase (SA-β-gal) staining

A549 and HSAEC were plated into 48 well-plates at density of 10,000 cells/well and 16,000 cells/well respectively. Cells were transfected 24 hours prior to bleomycin challenge at mRNA doses previously specified. Bleomycin (low dose: 0.1 µg/mL, high dose: 1 µg/mL) was applied to cells for 48 hours and was removed followed by 3x PBS washes. SA-β-gal activity was assessed at days 8 and 16 post-bleomycin exposure per manufacture’s protocol (Cell Signaling, #9860). Briefly, cells were washed with PBS and fixed in 1x fixative solution for 15 minutes. Staining solution containing X-gal diluted in dimethylformamide was generated and added to cells overnight at 37° C in a dry incubator. RBG brightfield images were obtained using a brightfield microscope. Percent positive cells were identified and quantified in Image J. >50 cells were analyzed per condition, across 3 biologically independent replicates.

### Air-liquid interface (ALI) cultures

Primary HSAECs were differentiated at the air-liquid-interface using an established manufacture’s protocol (STEMCELL Technologies). Briefly. HSAECs were expanded in submerged culture in PneumaCult-Ex Plus Medium (STEMCELL Technologies, #05040), prepared by supplementing PneumaCult-Ex Plus Basal Medium with 50X Supplement and hydrocortisone stock solution. Cells at passage <4 were seeded onto 12 mm Transwell inserts (STEMCELL Technologies, #05021) at 1.1 × 10⁵ cells per insert. 0.5 mL PneumaCult™-Ex Plus Medium was added apically, with 1 mL in the basolateral chamber. Medium was replaced in both chambers every other day until cultures reached full confluence. Cultures were then air-lifted by removing apical medium, and the basolateral chamber was changed to complete PneumaCult™-ALI-S Medium (STEMCELL Technologies, #05050), prepared from ALI-S Basal Medium with 10X Supplement and 100X Maintenance Supplement. Basolateral medium was replaced every 2 days, and apical surfaces were washed with pre-warmed PBS weekly to remove accumulated mucus. Cultures were differentiated for 28 days prior to use.

Differentiated human donor bronchial epithelium and pig airway ethmoid epithelium ALI cultures were provided by the University of Iowa *In Vitro* Models and Cell Culture Core. These were generated as previously described (*33*). Briefly, after enzymatic disassociation of bronchus and ethmoid epithelia, cells were seeded onto collagen-coated polycarbonate Transwell inserts (Corning Costar). ALIs were submerged in Ultroser G (USG) medium for 24 hours (37°C and 5% CO2). After 24 hours, the apical media was removed to promote differentiation at an air-liquid interface.

Fully differentiated ALI cultures were transfected apically with FLuc mRNA-LNPs at 10 µg mRNA for 24 hours. Luminescence was measured 24 hours post-transfection using an established luciferase assay system (Promega, E1500). Luminescence was assessed using the plate reader function on the IVIS imaging system.

### *In vivo* biodistribution

All animal procedures were approved by the University of Iowa IACUC (protocol #5022416). 10 week old CD-1 female mice received intratracheal instillation of post-nebulized saline or FLuc mRNA-LNPs (KC2/PEG formulation as previously specified) at a total mRNA dose of 12.5 µg. At 6 hours post-instillation, animals received an intraperitoneal 150 mg/kg dose of D-Luciferin. 10 minutes after D-Luciferin injection, animals were euthanized and kidney, spleen, liver, and lung were harvested and imaged ex vivo using an IVIS imaging system. Total flux was quantified per organ using the IVIS imaging software. Left and right organ flux for lungs and kidneys were added together to obtain the total organ flux per pair.

### *Ex vivo* sheep lung slices

Cheviot sheep lung slices were obtained from Ovis Sheep. Lung slices were maintained in DMEM with 10% FBS at 4° C during transportation from orthopedics surgical suite to the lab. Upon arrival to the lab, 1-3mm slices were obtained from lung tissue and placed into 37° C DMEM with 10% FBS. FLuc mRNA-LNPs at a dose of 10 µg mRNA or saline control were added to lung tissue media. 6 hours post-transfection, lung slices were submerged in 15mg/mL solution of D-Luciferin for 10 minutes and bioluminescence was subsequently measured using the IVIS system. Total lung slice flux was quantified per slice using the IVIS imaging software.

### Statistical analysis

All analyses were performed in GraphPad Prism. Data are presented as mean ± SEM unless otherwise noted. Sample sizes are stated in each figure legend. All tests were two-tailed, and a p-value < 0.05 was considered significant (*p < 0.05, **p < 0.01, ***p < 0.001, ****p < 0.0001; ns, not significant).

#### LNP physicochemical characterization (*Figure 2B*)

Z-average hydrodynamic diameter, polydispersity index, zeta potential, and encapsulation efficiency were measured on the same LNP preparations before and after nebulization and compared by Welch’s unpaired t-test (n = 3 independent preparations).

#### In vivo biodistribution (*Figure 2C*)

Organ total flux values were log₁₀-transformed and compared by two-way ANOVA with treatment and organ as factors, followed by Šídák’s multiple comparisons test comparing saline versus FLuc mRNA LNP within each organ (n = 5 mice per group).

#### ALI culture and ex vivo lung transfection (*Figure 2D, 2E*)

Luminescence values were log-10 transformed and compared between untransfected and FLuc mRNA LNP conditions by Welch’s unpaired t-test (n = 3 biological replicates per model). Exact p values are reported for non-significant comparisons.

#### Expression kinetics and proliferation (*Figure 3B-3E*)

Mean fluorescence intensity and cell counts are presented descriptively as mean ± SEM of 3 biological replicates; no hypothesis tests were performed on these curves.

#### γH2AX foci and alkaline comet assay (*Figure 4*)

Forty cells were scored per condition in each of 3 independent experiments. Differences in foci per cell were determined with one-way ANOVA with Tukey’s post hoc test. Individual cell values are displayed in the figures.

#### Senescence-associated β-galactosidase (*Figure 5B, 5C*)

The percentage of SA-β-gal-positive cells in GFP- and Dsup-pretreated bleomycin-exposed groups were compared at days 8 and 16 by two-tailed paired t-tests, matched by independent experiment (n = 3 biological replicates).

## Acknowledgement

We would like thank the team at SayoStudio for their illustration in Figure 1. The authors also thank the staff of the University of Iowa Central Microscopy Research Facility (CMRF) for training and technical guidance in image acquisition on the Zeiss 990 Confocal Microscope. We would also like to thank the University of Iowa Cell and Tissue Culture Core for providing cultured airway epithelial cells for this project. Lastly, we would like to thank the orthopedics team for providing fresh sheep lung tissue.

## Data Availability Statement

The data that support the findings of this study are available from the corresponding author upon reasonable request.

## Declaration of AI Use

During the preparation of this work, we used Claude (Anthropic) to assist with editing the text. After using this tool, we reviewed and edited the content as needed and take full responsibility for the content of the published article.

